# *In silico* Elucidation of Dihydroquinine Mechanism of Action against *Toxoplasma gondii*

**DOI:** 10.1101/2021.12.30.474617

**Authors:** Joseph A. Ayariga, Aarin M. Huffman, Audrey Napier, BK Robertson, Daniel A. Abugri

## Abstract

Dihydroquinine (DHQ), is a quinine-based compound with anti-malarial properties. However, little is known about its mechanism of action against *T. gondii* inhibition, which shares similar biology with *Plasmodium* spp. In order to explore DHQ activity as an inhibitor of *T. gondii* using *in vitro* assays, we first used an *in silico* approach to decipher its mechanisms of action based on previous knowledge about its disruption of nucleic acid and protein synthesis. An *in silico* study was performed on T. gondii parasite replication, transcriptional and translational machinery to decipher the binding potentials of DHQ to some top selected enzymes. We report for the first time, using an *in silico* analysis that showed that DHQ binds strongly to DNA gyrase, Calcium Dependent Protein Kinase 1 (CDPK 1), and prolyl tRNA synthetase and thus could affect DNA replication, transcriptional and translational activities in *T. gondii*. Also, we found DHQ to effectively bind to mitochondria detoxifying enzymes (i.e., superoxide dismutase (SOD), peroxidoxin, and Catalase (CAT)). In conclusion, DHQ could be a lead compound for the treatment of toxoplasmosis when successfully evaluated using *in vitro* and *in vivo* models to confirm its effectiveness and safety.

## Introduction

Zoonotic parasitic infections continue to cause serious public health, veterinary and socioeconomic predicament globally (Ben-Harari and Connolly, 2019; Pappas et al., 2009; Flegr et al., 2014; Dubey, 2010). For example, toxoplasmosis, the disease caused by *T. gondii* affects nearly 1/3 of the human population worldwide (Pappas et al., 2009; Flegr et al., 2014). More worrisome, a recent study, revealed that over 35 to 76% of wild and domestic felids are infected with *T. gondii* (Montazeri et al., 2020). This suggests that its infection in man and animals are on the increase globally. To overcome these parasitic diseases, different strategies are required which include, management, prevention, and treatment with safe and effective chemical inhibitors. Although few drugs are available for the treatment of individuals (humans and animals) infected with *T*.*gondii*, they are faced with limitations such as toxicity, high cost, and more so, most drugs are ineffective in treating the latent stage (bradyzoite) that continuously persist in the brain (Shammaa et al., 2021; Secrieru et al., 2020; Angel et al., 2020; Shiojiri et al., 2019). Thus, there is an urgent call for the development of novel compounds and inhibitors against this parasite, and more importantly an in-depth analysis and identification of the specific targets for safe and effective druggability and treatment.

DHQ is a pharmaceutical impurity associated with quine production (Nontprasert et al., 1996). It has been reported to have anti-malarial properties (Gorka et al., 2013; Achan et al., 2011; Sanchez et al., 2008; Nontprasert et al., 1996; Pukrittayakamee et al., 1997; Brossi et al., 1973; Fayer et al., 1972; Brossi et al., 1971; Polet and Barr, 1968). Furthermore, its mechanism of action has been predicted in *Plasmodium* species to target nucleic acid and protein synthesis (Brossi et al., 1973; Brossi et al., 1971). However, its mechanism of action against *T. gondii*, which shares a similar phylum with *Plasmodium* is yet to be investigated.

Here, using computational analysis, we seek to assess the molecular mechanism of action of DHQ in *T. gondii* using selected replicative, transcriptional, translational, and mitochondria machinery such as DNA gyrase, CDPKs, MIF, RNA synthase, SOD, peroxidoxin, and Catalase.

## Author Summary

Early deciphering of compounds’ mechanism of action is crucial for drug discovery and development. This approach saves time, resources and provides an insight into the possible mechanism of action of compounds of interest before wet experimental work could be carried out. In this paper, we used *in silico* approach as a first point of deciphering the mechanism of DHQ, a quinine derivative, that has been reported to have anti-*Plasmodium* activity *in vitro* and *in vivo*. The prediction showed that DHQ binds strongly to very important replicative, transcriptional, translational, and mitochondrial-associated proteins. Thus, this process was carried out as the first approach to predict DHQ’s possible mechanism of action against *T. gondii* before performing wet labs to gain insight into the compound’s inhibitory activity in *T. gondii* growth *in vitro*.

## Results and Discussion

### *In silico* Docking of DHQ with *T. gondii* Replicative and Translational Machinery

Based on our cellular and biochemical assay results, we used *in silico* docking analysis to decipher whether DHQ could be targeting the following enzymes/receptors: DNA Gyrase, TgCDPK1, tRNAs, MMIF, ROP5B, and ROP5C. These proteins play crucial roles in the parasite’s invasion, survival, and replication. Herein, we analyzed for the theoretical binding affinities between DHQ and the enzymes/receptors as well as their ligand-receptor interactions at the docking pocket. The docking analysis was performed using extracted crystalized structures of the receptors from the RCSB website or modeled using the online modeling software SWISS-MODEL (https://swissmodel.expasy.org/). DHQ, 3-Amino-1H-pyrazole-4-carboxamide, and Moxifloxacin were downloaded from the PubChem database (https://pubchem.ncbi.nlm.nih.gov/).

### DHQ interaction with *T. gondii* DNA Gyrase interactions

To stop the progression of multidrug-resistant tuberculosis, fluoroquinolone has been shown to have high potency, and the mechanism of action of fluoroquinolone is known to target DNA gyrase (Blower et al., 2016). It has also been proven in parasites (e.g., Trypanosomes) that some quinolone drugs bind to DNA gyrase to form a complex that blocks transcription by RNA polymerase (Willmott et al., 1994). DNA gyrase of *T. gondii* has also been indicated as a potential target for drugs against *T. gondii* (Lin et al., 2015). While the crystal structure of the DNA gyrase of multidrug-resistant tuberculosis has been discovered, the DNA gyrase of *T. gondii* is currently unavailable. In this work, we used the crystal structure of multidrug-resistant tuberculosis as a model study receptor to dock DHQ as a ligand and to analyze for their binding affinities (ΔG). To also show if there exist any contrasting binding affinities between DHQ and fluoroquinolone for the *M. tuberculosis* DNA gyrase as well as *T. gondii* DNA gyrase, their binding affinities were compared and the specific amino acids interacting with these compounds were analyzed for the type of interactions occurring at the binding pocket. While fluoroquinolone (also known as moxifloxacin) showed a slightly lower binding affinity of -7.2 kcal/mol (**Table 1**) to gyrase of *M. tuberculosis*, DHQ bonded to *T. gondii* (TgDNA) gyrase with a binding affinity of -8.0 kcal/mol (**Table 1**) compared to DHQ - *M. tuberculosis* DNA gyrase interaction which produced a relative free binding energy of -6.6 kcal/mol (**Table 1, Figure 2A, 2B and 2E**). As shown in **Figure 1**, the 3D rendition of DHQ in red color interacting with gyrase in green color, also in **Figure 1**, a 2D depiction of DHQ interacting with gyrase at the binding pocket indicates that DHQ forms a strong hydrogen bond with ARG234 producing a bond distance of 1.90Å, for which DHQ acts as the hydrogen acceptor and ARG234 the donor. The ALA255 of TgDNA gyrase binds to DHQ via hydrophobic interactions, forming a pi-alkyl bond with DHQ with a bond distance of 4.69 Å in the binding pocket. Forming a hydrogen bond at 2.97Å with DHQ, TgDNA gyrase via ARG234 acts as a hydrogen donor for which DHQ is the acceptor. DHQ was able to form a stable hydrogen bond with TgDNA gyrase via the gyrase’s ASP200 amino acid residue which acts as a hydrogen acceptor. A bond distance of 1.99 Å is established between them. ARG195 interacts with DHQ via hydrogen bond also, for which ARG195 is the hydrogen acceptor. LEU257 of TgDNA gyrase establishes pi-sigma interaction with DHQ to establish a hydrophobic bond of distance 3.38 Å. Other hydrophobic bonds formed by TgDNA gyrase with DHQ established in the binding pocket includes amino acids ALA203 (bond distance, 3.55 Å), ALA255 (bond distance, 5.26 Å), LEU257 (bond distance, 5.32 Å), ALA267 (bond distance, 4.72 Å), LEU230 (bond distance, 4.94 Å), LEU257 (bond distance, 4.62 Å) and ALA255 (bond distance, 4.69 Å). This analysis points to stable DHQ-DNA gyrase interactions, illustrating the possible mechanism of action of DHQ as a *T. gondii* inhibitor via the inhibition of DNA replication through the binding to and interfering in gyrase activity during the parasite’s DNA replication. This knowledge hints at a possible pharmacological utility of DHQ against *T. gondii* and multidrug-resistant tuberculosis.

**Figure 1.**
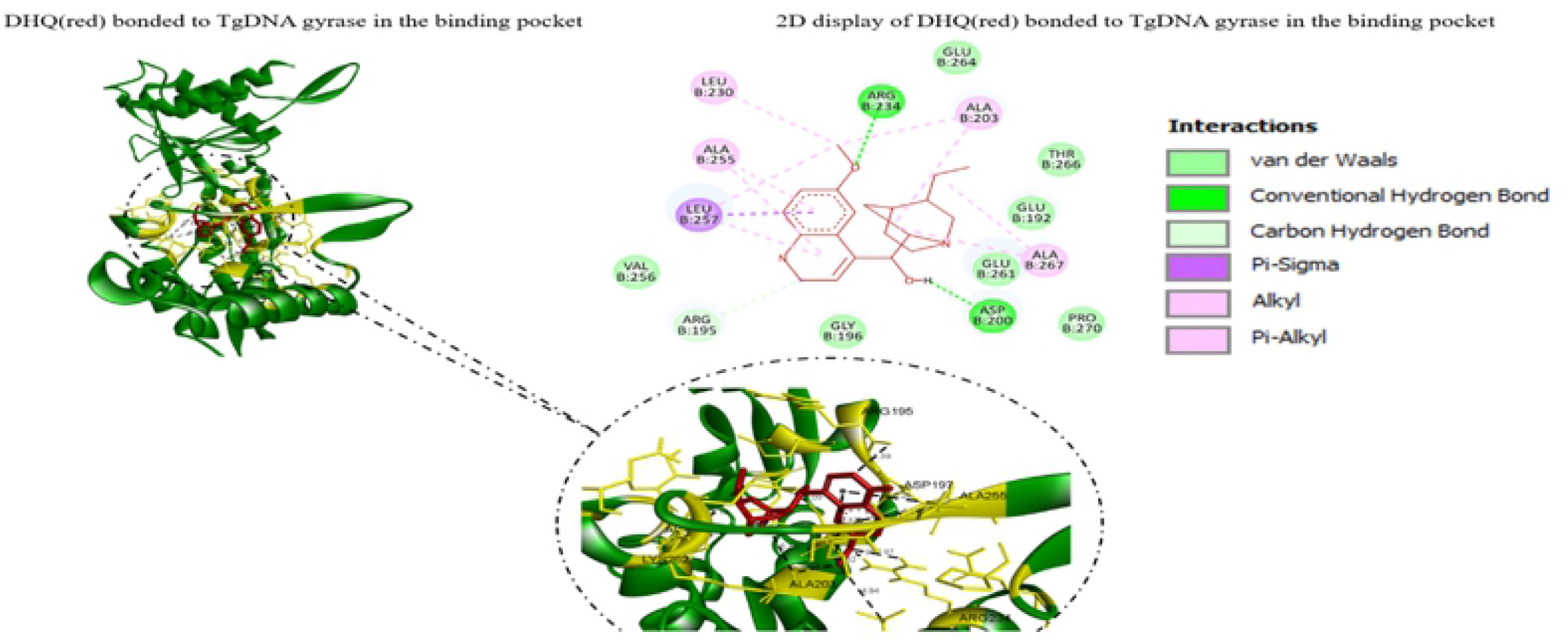
As shown in **Figure 1**, DI IQ forms a strong hydrogen bond with ARG234 producing a bond distance of I.90A. for which DHQ acts as the hydrogen acceptor and ARG234 the donor. The ALA255 of Tg DNA gyrase binds to DHQ via hydrophobic interactions, forming a pi-alkyl bond with DHQ with a bond distance of 4.69 Å in the binding pocket. Forming hydrogen bond at 2.97 Å with DI IQ. TgDNA gyrase via ARG234 acts as hydrogen donor for which DHQ is the acceptor. DHQ is able to form a stable hydrogen bond with TgDN/X gyrase via the gyrase’s ASP200 amino acid residue which acts as a hydrogen acceptor. A bond distance of 1.99 Å is established between them. ARG 195 interacts with DHQ via hydrogen bond also, for which ARG 195 is the hydrogen acceptor. LEU257 of TgDNA gyrase establishes pi-sigma interaction with DHQ to establish a hydrophobic bond of distance 3.38 Å. Other hydrophobic bonds formed by TgDNA gyrase with DHQ established in the binding pocket includes amino acids ALA203 (bond distance, 3.55 A). ALA255 (bond distance. 5.26 Å), LEU257 (bond distance. 5.32 Å), ALA267 (bond distance. 4.72 Å). LEU230 (bond distance. 4.94 Å). LEU257 (bond distance. 4.62 A) and ALA255 (bond distance. 4.69 Å).

**Figure 2.**
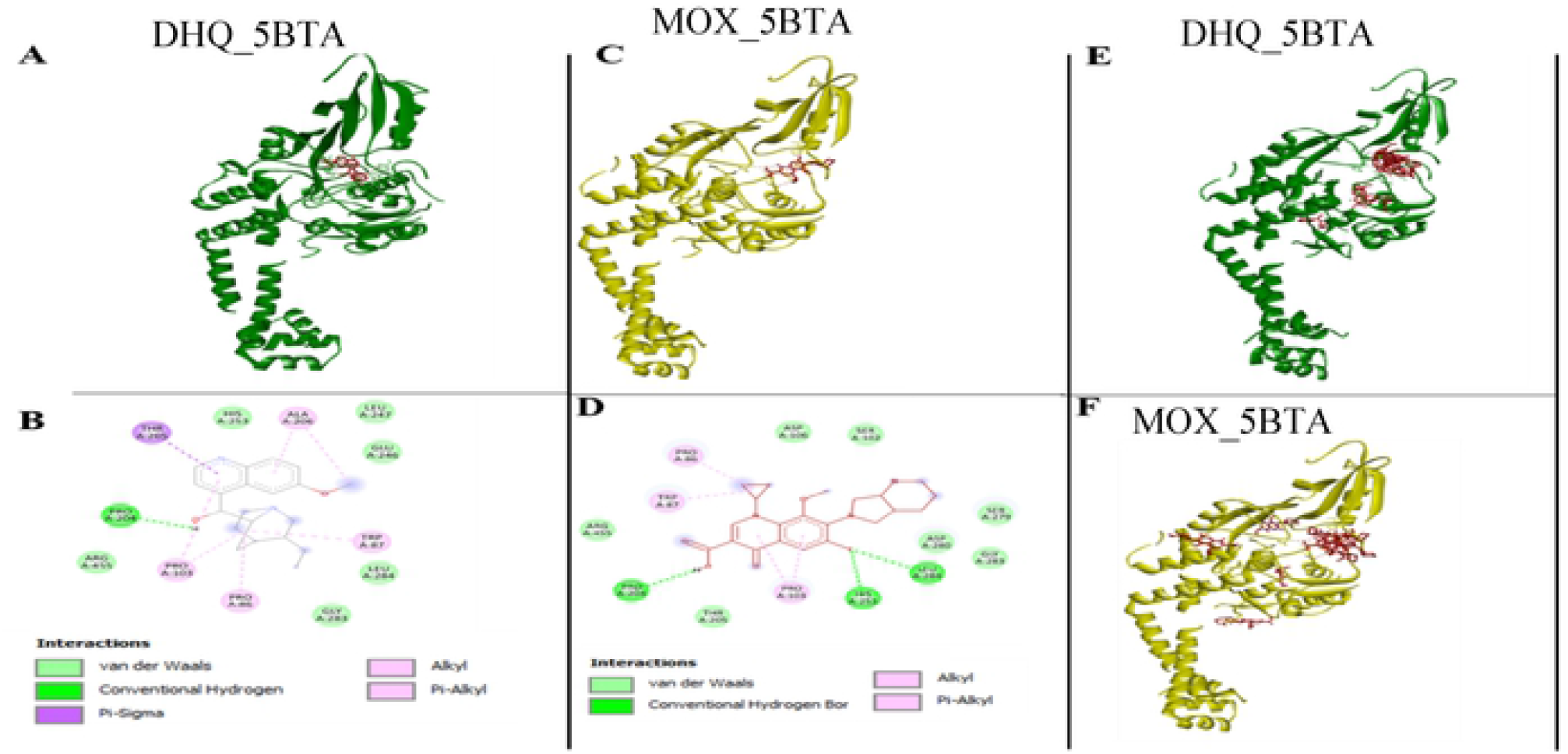
DNA gyrase A of *Mycobacterium tuberculosis*. **Figure 2A, 2B, 2E** shows DHQ interaction with gyrase A (5BTA)(a topoisomerase II complex of *Mycobacterium tuberculosis* H37Rv). As shown in Figure **2A**, the 3D rendition of DHQ in red color interacting with gyrase A in green color. Figure **2D** is a 2D depiction of DHQ interacting with gyrase at the binding pocket indicates that proline at amino acid position 204 interacts directly with DHQ via hydrogen bonding, whereas PRO86 and PRO 103 shows alkyl interaction with the compound. TRP87, and ALA206 also shows alkyl and pi-alkyl interaction with DHQ. A single pi-sigma bond is formed between THR205 and DHQ. **Figure 2C** depicts Moxifloxacin (red) bound to gyrase A (yellow), whereas Figure 2D shows a 2D rendition of moxifloxacin (red) interacting with the binding pocket atoms of gyrase A via 3 hydrogen bonds and four alkyl and pi-alkyl bonds. **Figures 2E** and **2F** also shows all the 9 different conformational poses of the DHQ (in red) bonding to gyrase A molecule in green and the 9 different poses of moxifloxacin (red) in the binding pockets of gyrase (yellow) respectively.

### DHQ and *T. gondii* prolyl tRNA synthetase interactions

Prolyl-tRNA synthetase belongs to the aminoacyl-tRNA synthetase family and plays a crucial role in protein translation in living cells. The tRNA synthetase of *T. gondii* has been a target for several compounds used as therapeutics against *T. gondii* and other Apicomplexans, e.g., the use of febrifugine and halofuginone (Mishra et al., 2019). This current work revealed that DHQ targets the prolyl tRNA synthetase of *T. gondii* with a high relative free binding energy of -7.7 kcal/mol (**Table 1**). **Figure 3C** is the 3D depiction of the DHQ interacting with the tRNA synthetase of *T. gondii*. Also illustrated in **Figure 3C and Figure 3D)**, a 2D illustration of the contacts of DHQ at the binding site to tRNA synthetase of *T. gondii* indicates that DHQ shares two hydrogen bonds at amino acids ARG594, and GLN555. There exists also a van der Waals interaction between the compound and the amino acid HIS560, a pi-alkyl interaction at amino acid PHE415 and pi-cation interaction with ARG470.

**Figure 3.**
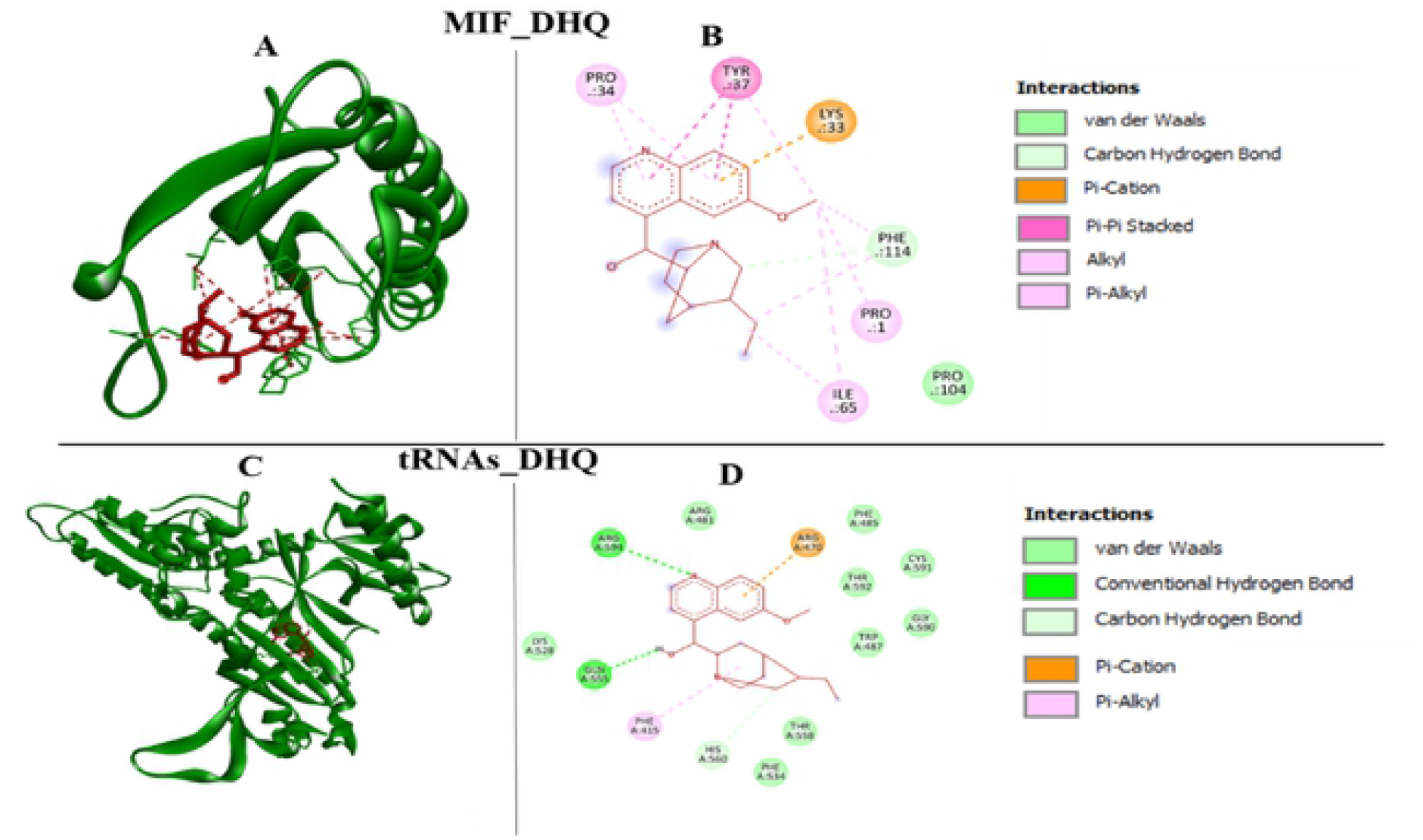
**Figure 3** shows DHQ binds stably to *T. gondii* MIF. As shown in **Figure 3A**, DHQ is shown bound to two loops of MIF molecule. The 2D rendering (**Figure 3B**) shows that the most stable conformation was achieved by the binding of DHQ to the PHEI14. PRO1. ILE65, LYS33.TYR37. and PRO34 via hydrogen bonding, alkyl, pi-alkyl, pi-cation, pi-pi stacked, and alkyl interactions respectively. **Figure 3C** is the 3D depiction of the DHQ interacting with tRNA synthetase of *T. gondii*. Whereas **Figure 3D**. a 2D rendition shows that DHQ binds to tRNA synthetase of *T. gondii* via two hydrogen bonds at amino acids ARG594, and GI.N555. There exists also a van der Waals force between the compound and the amino acid HIS560, a pi-alkyl interaction at amino acid PHE4I5 and pi-cation interaction with ARG470.

This analysis shows that there is a stable DHQ-prolyl tRNA synthetase interactions, pointing to the capacity of DHQ as *a T. gondii* inhibitor by interrupting the *T. gondii* protein translational activity by binding to the parasite’s prolyl tRNA synthetase.

### DHQ and Macrophage Migration Inhibitory Factor of *T. gondii* (MIF) Interactions

MIF is a proinflammatory molecule in mammals with homologs in parasites that possesses tautomerase and oxidoreductase enzymatic activities (Sommerville et al., 2013). Here we downloaded the tautomerase active MIF of *T. gondii* and studied the binding of DHQ to this molecule. Even though similar structural homology of *T. gondii* and mammalian MIFs do exist, their respective tautomerase active sites are different, and this justifies the variation in inhibitor sensitivity (Sommerville et al., 2013). We showed that *T. gondii* MIF binds stably to DHQ at a binding affinity of -6.9 kcal/mol **(Figure 3A)**. As shown in **(Figure 3A)**, DHQ is bound to two loops of the MIF molecule. The 2D rendering **(Figure 3B)** shows that the most stable conformation was achieved by the binding of DHQ to the PHE114, PRO1, ILE65, LYS33, TYR37, and PRO34 via hydrogen bonding, alkyl, pi-alkyl, pi-cation, pi-pi stacked, and alkyl interactions, respectively. Since MIF of *T. gondii* is known to play an immunomodulatory role during *the T. gondii* infection process (Sommerville et al., 2013), this stable binding of DHQ to MIF interferes and deregulates the parasite’s MIF proper function.

### DHQ and ROP5B/C (*T. gondii* virulent allele) pseudokinase interactions

ROP5B/C belongs to the ROP5 family which is a set of polymorphic pseudokinases. They have been demonstrated to play major roles in *T. gondii* pathogenesis (Reese et al., 2011; Reese et al., 2014). They play critical functions in subcellular localization of specific receptors, recognition of partner molecules, and regulators of signaling networks (Boudeau et al., 2006). In this study, we investigated the possibility of DHQ dysregulation of ROP5B by the compound binding to the pseudokinase and hindering its proper functional conformations. Our docking analysis showed that ROP5B interacted with DHQ with a high binding affinity of -6.9 kcal/mol. In **(Figure 4A)**, DHQ in red color is shown to bind to ROP5B (green).

**Figure 4.**
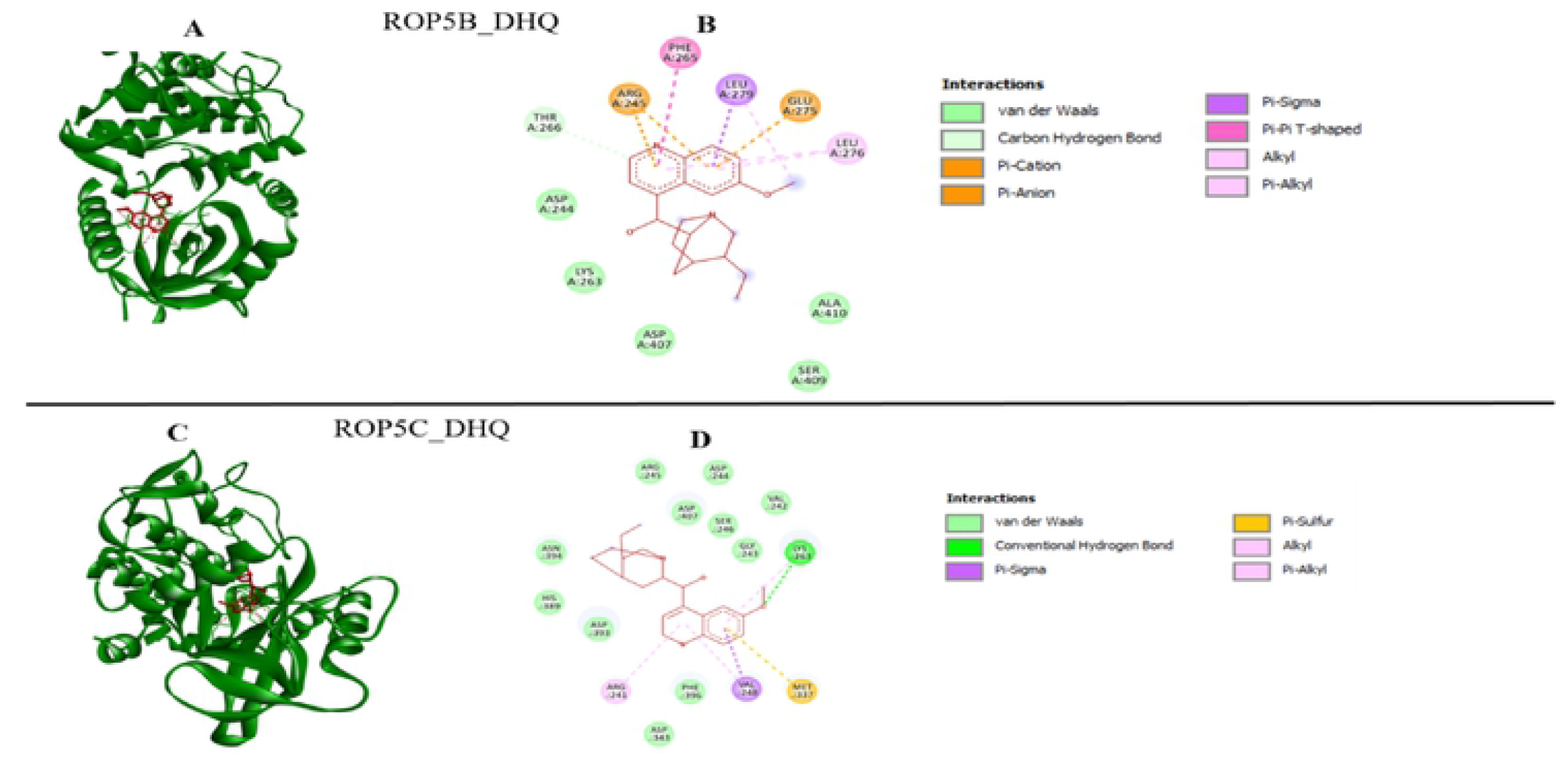
In **Figure 4A**, DHQ in red color is shown to bind to ROP5B (green). Also, the 2D depiction (**Figure 4B**) of the contacts of DHQ at the binding site to ROP5B pseudokinase of *T. gondii* shows DHQ interacting with amino acids ARG245. and GLU275 via pi-catioivanion interactions. DHQ established pi-pi-T shaped interaction with the pseudokinase at PHE265. At LEU279, there is a pi-sigma bond with the compound also. DHQ (in red) as shown in **Figure 4C** interact with ROP5C, and in **Figure 4D**. the 2D diagram shows a stable interaction at the binding pocket in which LYS263of ROP5C makes hydrogen bond contact with the compound. VAL248 makes a pi-sigma contact, whereas ARG24I makes an alkyl contact with DHQ. MET337 however makes a pi-sulfur bond with DHQ. Collectively, these interactions create a stable and strong bonding between DHQ and ROP5C, pointing to the capability of DI IQ as *T. gondii* inhibitor by the mechanism of the disruption of the *T. gondii* pathogenesis.

Also, the 2D depiction **(Figure 4B)** of the contacts of DHQ at the binding site to ROP5B pseudokinase of *T. gondii* shows DHQ interacting with amino acids ARG245, and GLU275 via pi-cation/anion interactions. DHQ established pi-pi-T shaped interaction with the pseudokinase at PHE265. Also, at LEU279, there was a pi-sigma bond with the compound. The ROP5C pseudokinase (Reese et al., 2011), on the other hand, showed a binding affinity of -7.4 kcal/mol (**Table 1**) with DHQ. DHQ is shown in **Figure 4C** to interact with ROP5C, and in **Figure 4D**, the 2D diagram shows a stable interaction at the binding pocket in which LYS263 of ROP5C makes hydrogen bond contact with the compound, VAL248 makes a pi-sigma contact, whereas ARG241 makes an alkyl contact with DHQ. MET337 however, makes a pi-sulfur bond with DHQ. Collectively, these interactions create stable and strong bonds between DHQ and ROP5C, pointing to the capability of DHQ as a *T. gondii* inhibitor by the mechanism of binding to and disrupting the *T. gondii* pathogenesis process.

### DHQ and Calcium-Dependent Protein Kinase 1 from *T. gondii* (TgCDPK1) interactions

The CDPK1 enzyme in *T. gondii* controls multiple processes that are crucial for the parasite’s intracellular replicative cycle and the parasite’s secretion of adhesins (Ojo et al., 2010; Johnson et al., 2012). This enzyme plays a major function in the ability of the parasite to invade host cells, move, and egress (Fluorentine et al., 2017). We analyzed how effective DHQ binds to TgCDPK1 and thus affects the replicative and invasion properties of the parasite. Interestingly, the binding affinity of DHQ to TgCDPK1 was -6.7 kcal/mol, a -1.2 kcal/mol higher than a known inhibitor of TgCDPK1 called 3-Amino-1H-pyrazole-4-carboxamide which recorded -5.5 kcal/mol. As shown in (**Figure 5A, and Figure 5B**) the ligand-receptor interaction showed that DHQ interacts with TgCDPK1 via two hydrogen bonds emanating from LEU345 and ARG439, two alkyl bonds formed between Lys59, LYS435, and LEU438, and finally a pi-anion interaction between GLU64 and DHQ. The -5.5 kcal/mol of 3-Amino-1H-pyrazole-4-carboxamide interaction with TgCDPK1 came from the binding of the compound to the active site with the following residues: ILE212, LYS338, GLN334, GLN341, and GLU178 which all demonstrated conventional hydrogen bonding with the compound except GLU178 which established a pi-anion bond with 3-Amino-1H-pyrazole-4-carboxamide **(Figure 5C, and 5D)**.

**Figure 5.**
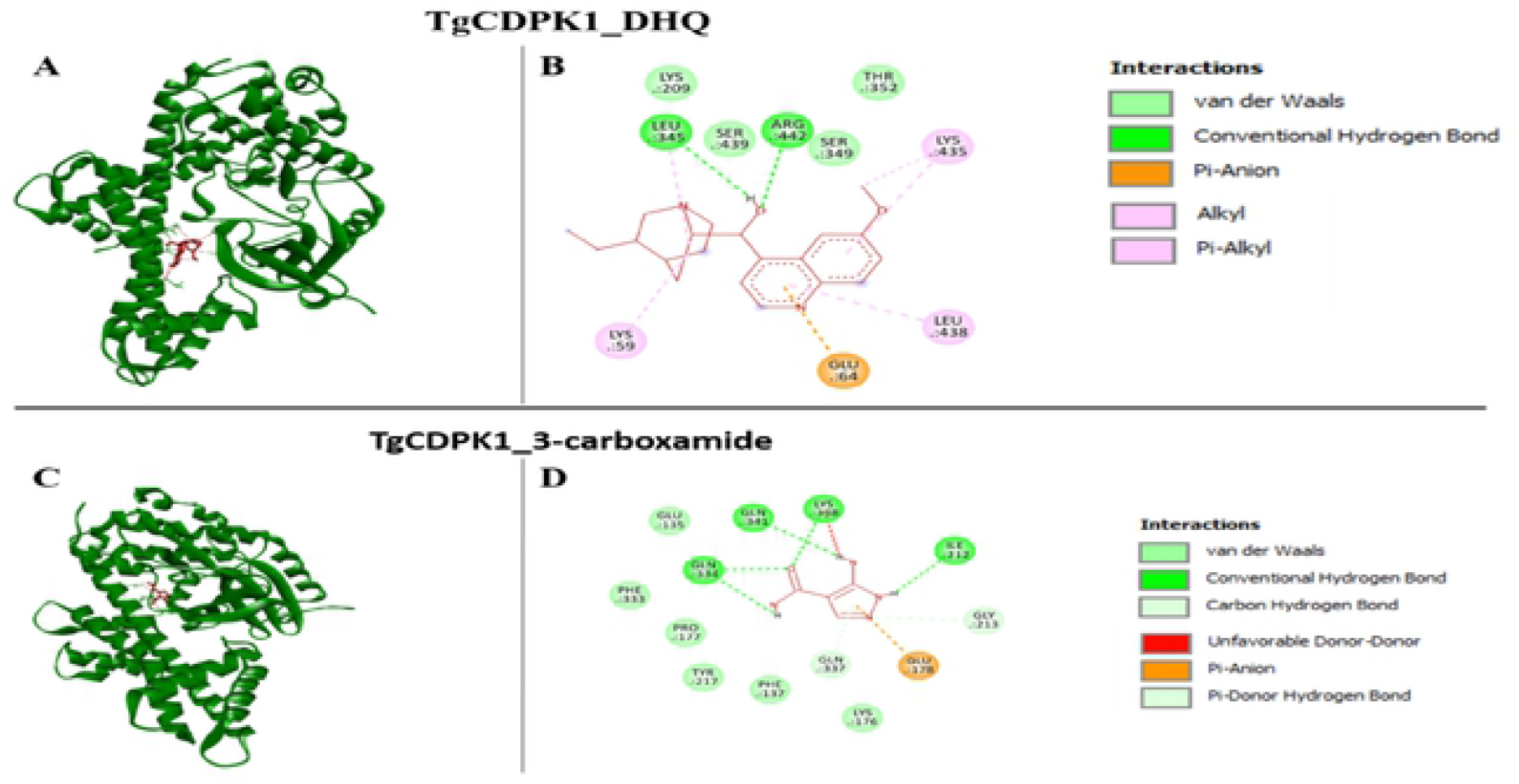
**Figure 5A** shows the 3D rendition of DHQ-TgCDPKI interaction. **Figure 5B** shows a 2D diagrammatic illustration of DHQ interacting with TgCDPKl via two hydrogen bonds emanating from LEU345 and ARG439, two alkyl bonds formed between Lys59, LYS435 and LEU438 and finally a pi-anion interaction between GLU64 and DHQ. In **Figure 5C**, 3-Ainino-l H-pyrazole-4-carboxamide is depicted interacting with TgCDPKl, and **Figure 5D** shows the 2D rendition of the interaction which emanates from the binding of the compound to the active site of TgCDPKl at the following residues; ILE212, LYS338.GLN334. GLN34I, and GLUI78 which all demonstrated conventional hydrogen bonding with the compound except GLUI78 which established a pi-anion bond with 3-Ainino-l H-pyrazole-4-carboxamide. LYS338 however shows a strong unfavorable donor-donor interaction with the hydrogen moiety of 3-Amino-lH-pyrazole-4-carboxamide. This interaction therefore caused a considerable loss of affinity of 3-Ainino-1H-pyrazole-4-carboxamide to TgCDPK I compared to DHQ.

LYS338 however, shows a strong unfavorable donor-donor interaction with the hydrogen moiety of 3-Amino-1H-pyrazole-4-carboxamide. This interaction therefore might have caused a considerable loss of affinity of 3-Amino-1H-pyrazole-4-carboxamide to TgCDPK1 compared to DHQ. Hence, the strong unfavorable donor-donor interaction recorded seems to slightly dampen this stability of 3-Amino-1H-pyrazole-4-carboxamide to TgCDPK1 interaction, hence the recorded -5.5 kcal/mol binding affinity, DHQ on the other hand which produced a very high affinity of -8 kcal/mol recorded no unfavorable bonds in its binding pocket. Hence, we predict a higher inhibitory potential for DHQ on TgCDPK1 than 3-Amino-1H-pyrazole-4-carboxamide. The binding of DHQ or 3-Amino-1H-pyrazole-4-carboxamide to TgCDPK1 prevents *T. gondii* phosphorylation and ATP binding activities (Ravi et al., 2020).

### DHQ interaction with *T. gondii* Mitochondria proteins (SODs, Peroxiredoxins, and Catalase)

Mitochondria is a well-known important energy organelle in eukaryotes and especially, apicomplexans parasites e.g., *T. gondii*. The organelle’s basic functions include cytosolic nucleic iron-sulfur proteins detoxification, nucleic acid synthesis, fatty acid synthesis, and protein synthesis as well as generation of energy for cellular mechanistic activities (Syn et al., 2017; Lill et al., 2014; Kwok et al., 2004; Melo et al., 2000). Thus, cellular activities that cause mitochondria imbalances could lead to disruption of the above functions mentioned as well as the membrane potential in *T. gondii*.

For instance, the common causes of mitochondria imbalances in the cell could be partly due to the induction of external and internal stimuli such as stress, chemical inducers, and microbial infections (Syn et al., 2017; Kwok et al., 2004; Callahan et al., 1988; McGonigle and Dalton,1998; Nathan and Shiloh, 2000). The inducers of mitochondria function often cause the excessive generation of reactive oxygen species (ROS) (e.g., hydrogen peroxide, superoxide radical, and hydroxyl radical) by parasites (Kwok et al., 2004; Vercesi et al., 1998). It has been well documented that this ROS creates microenvironmental oxidative stress intracellularly which causes toxicity to parasites and eventually parasites death (Kwok et al., 2004). To overcome this ROS toxicity in eukaryotic and especially parasites cells, the mitochondria use its internal enzymes such as superoxide dismutase, catalase, glutathione peroxidase, and peroxiredoxins to form an antioxidant to detoxify the reactive oxygen species (ROS) (Kwok et al., 2004; Son et al., 2001; Odberg-Ferragut et al., 2000; Ding et al., 2000; Kaasch and Joiner, 2000; Vercesi et al., 1998; Miller and Britigan, 1997).

To validate whether DHQ had any way of counteracting the mitochondrial enzymes that aid *T. gondii* to detoxify drugs and other induces of ROS, we screen DHQ against the above-mentioned enzymes. Interestingly, DHQ interaction with *T. gondii* peroxidoxin (PRX), SOD3, and CAT had binding affinities of -7.0, -6.9, -6.8 kcal/mol, respectively. **Figure 6A** and **6B** depicts DHQ (red) bound to the SOD3 molecule in the binding pocket. As shown in **Figure 6B**, DHQ establishes a stable hydrogen bond with GLU210. It also interacts with SOD3 via alkyl and pi-alkyl hydrophobic interactions through amino acids HIS80, TYR84, PHE168, and ARG221. Other weaker van der Waals interactions established between SOD3 and DHQ were recorded on amino acids LYS166, GLY169, ASN87, THR165, HIS167, TRP 209, HIS211, ASN 219, ASP220, and GLY222. Together these interactions enhanced the stability of the binding of DHQ to the SOD3 molecule. Figures **6C** and **6D** depict the interaction of DHQ (red) with the SOD2 molecule. As shown in **Figure 6C**, DHQ formed a stable hydrogen bond with ALA118 and HIS115. It also interacts with SOD2 via alkyl and pi-alkyl hydrophobic interactions through amino acids LYS122, TYR119, ARG260, and PRO261. The weak van der Waals interactions established between SOD2 and DHQ were recorded on amino acids ASN151, ALA203, PHE206, GLY207, TRP248, GLU249, ASN258, and ASP259. Accumulatively these interactions boosted the stability of the biding of DHQ to SOD2 molecule.

**Figure 6.**
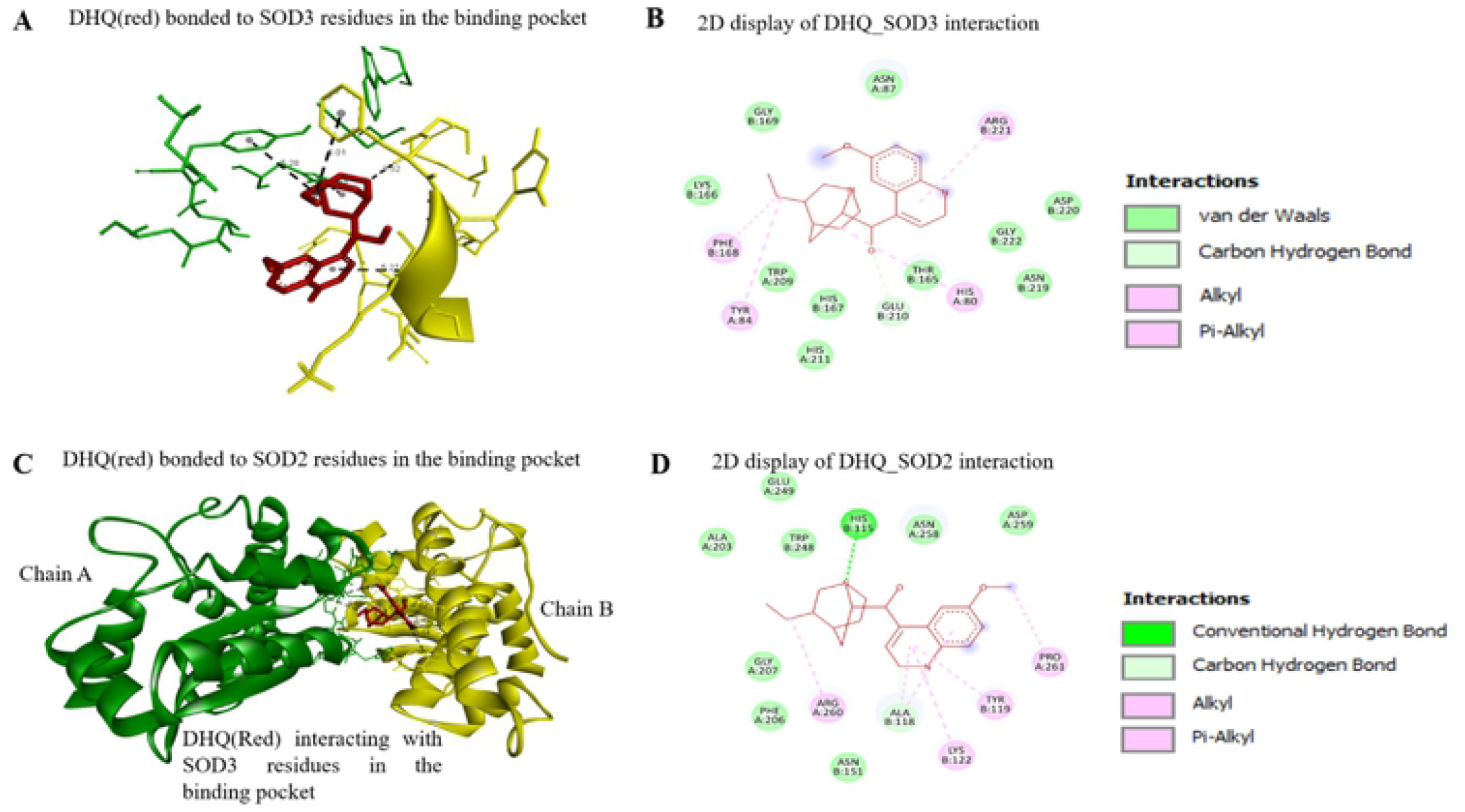
In **Figure 6A** and **6B** depicts DI IQ (red) bound to S0D3 molecule in the binding pocket. As shown in the **Figure 6B**, DHQ establishes stable hydrogen bond with GLU2I0. It also interact with SOD3 via alkyl and pi-alkyl hydrophobic interactions through amino acids HIS80, TYR84, PHE168, and ARG22I. Other weaker van der Waals interactions established between SOD3 and DHQ were recorded on amino acids LYS166, GLY169, ASN87, THR165, HIS167. TRP 209. IIIS21I. ASN 219. ASP220 and GLY222. Together these interactions enhanced the stability of the binding of DHQ to SOD3 molecule. The **Figure 6C** and **6D** depicts the interaction of DHQ (red) with SOD2 molecule. As shown in the **Figure 6C**. DI IQ formed a stable hydrogen bond with ALAI 18 and HIS 115. It also interact with SOD2 via alkyl and pi-alkyl hydrophobic interactions through amino acids LYS122, TYR119. ARG260. and PRO261. The weak van der Waals interactions established between SOD2 and DHQ were recorded on amino acids ASN151. ALA203. PHE206, GLY207. TRP248, GLU249, ASN258. and ASP259. Accumulatively these interactions boosted the stability of the binding of DHQ to SOD2 molecule.

Figure **7A** and **7B** depicts DHQ (red) bound to Tg Catalase chain A. As depicted in **Figure 7B**, DHQ forms a stable hydrogen bond with HIS183 and shares a pi-sigma bond with VAL291 and VAL443 respectively. VAL291 and VAL443 can also interact with DHQ via alkyl hydrophobic interactions. Tg Catalase also can bind to DHQ via its PRO293 by another hydrophobic alkyl interaction with the hydroxyl group of the DHQ. **Figures 7C** and **7D** depict the interaction of DHQ (red) with the Tg peroxidoxin molecule. As shown in **Figure 7D**, Tg peroxiredoxin binds tightly to DHQ via two hydrogen bonds established between DHQ at amino acids AS204 and TYR164. It is also able to stabilize DHQ in the binding pocket via hydrophobic interactions such as pi-pi-T shaped interactions, alkyl, and pi-alkyl interactions through the amino acids TYR164, IL3106, VAL206, LEU128, PHE132, LEU228, and PRO 127. Also, a weaker van der Waal force exists between ALA163, SER188, LEU190, THR131, and ALA269 with DHQ. These altogether stabilized DHQ in Tg peroxiredoxin binding pocket.

**Figure 7.**
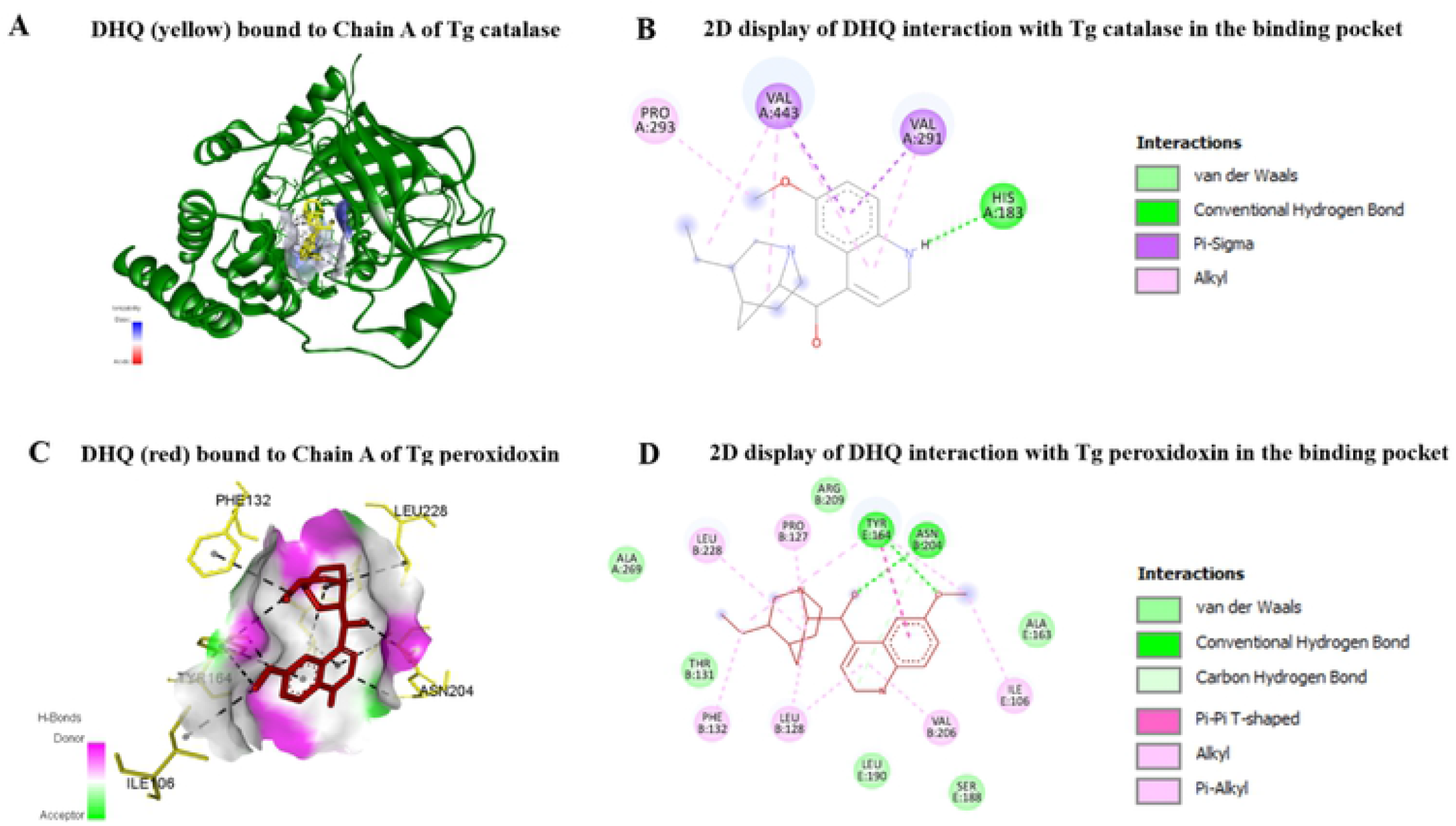
In **Figure 7A** and **7B** depicts DI IQ (red) bound to Tg Catalase chain A. As depicted in the **Figure 7B**. DHQ forms stable hydrogen bond with HIS 183, and shares a pi-sigma bond with VAL291 and VAL443 respectively. VAL291 and VAL443 are able to also interact with DHQ via alkyl hydrophobic interactions. Tg Catalase also is able to bind to DHQ via its PRO293 by another hydrophobic alkyl interaction with the hydroxyl group of the DHQ. The **Figure 7C** and **7D** depicts the interaction of DHQ (red) with Tg peroxidoxin molecule. As shown in the **Figure 7D**. Tg peroxidoxin binds tightly to DHQ via two hydrogen bonds established between DHQ at amino acids AS204 and TYR164. It is also able to stabilize DHQ in the binding pocket via hydrophobic interactions such as pi-pi-T shaped interactions, alkyl and pi-alkyl interactions through the amino acids TYRI64, 1L3106. VAL206, LEU 128. PHE132, LEU228 and PRO 127. Also, a weaker van der Waal force exist between ALA 163, SER 188. LEU 190. THR131 and ALA269 with DHQ. These altogether stabilized DHQ in Tg peroxidoxin binding pocket.

Our *in-silico* studies, corroborated with the mechanism(s) of action of DHQ previously reported in *Plasmodium* species to inhibit DNA replication, RNA, and protein synthesis (Achan et al., 2011; Nontprasert et al., 1996; Brossi et al., 1973; Brossi et al., 1971; Polet and Barr, 1968).

It has been well documented that in multidrug-resistant (MDR) tuberculosis treatment, quinolone-based drugs such as fluoroquinolone can effectively inhibit the bacteria by targeting its DNA gyrase (Blower et al., 2016). Another study in Trypanosomes has also elucidated that DNA gyrase perfectly binds with quinolone drugs thus forming a unique complex that blocks transcription at the RNA polymerase site (Willmott et al., 1994). Thus, DHQ might be targeting the DNA gyrase of *T. gondii* which has been identified as a potential target for drugs against *T. gondii* (Blower et al., 2016). Additionally, the observed effective and stable DHQ-prolyl tRNA synthetase interactions obtained from the docking studies point to the possibility of DHQ inhibiting *T. gondii* replication and invasion by interrupting the *T. gondii* protein translational activity by strongly binding to the parasite’s prolyl tRNA synthetase.

Calcium-dependent protein kinases are very important to *T. gondii* invasion and survival processes (Ojo et al., 2010; Johnson et al., 2012). Interestingly, our docking studies showed that DHQ or 3-Amino-1H-pyrazole-4-carboxamide effectively binds to TgCDPK1 and thus could avert its phosphorylation and ATP binding activities (Blower et al., 2016). The anti-proliferation and anti-invasion results obtained in the *in vitro* studies corroborated with our *in silico* docking studies that indicate that DHQ effectively binds to TgCDPKI. In previous studies, by (Ojo et al., 2010; Johnson et al., 2012), TgCDPKI has been discovered to play a crucial role in *T. gondii* proliferation and invasion of host cells. Thus, the strong binding affinity obtained implies that DHQ could block TgCDPKI expressions and hence blocks invasion and prevent the proliferation of *T. gondii*.

The *T. gondii* peroxidoxin (PRX), SOD3, and CAT’s strong interaction with DHQ observed, suggested that DHQ could be an excellent inhibitor of the *T. gondii* mitochondria ROS detoxifying enzymes. Thus, could cause parasites death due to excessive toxicity of the generated ROS caused by the immune-related cells to offset any microbial infection and chemical inducers such as drugs (Syn et al., 2017; Kwok et al., 2004; Callahan et al., 1988; McGonigle and Dalton,1998; Nathan and Shiloh, 2000). However, since this is *in silico*, further studies are needed to confirm this conjecture.

## Conclusion

DHQ was highly effective in binding to replicative, transcriptional, and translational machinery of *T. gondii in silico*. Thus, could cause inhibition of *T. gondii* growth, invasion, and egress. Also, based on the *in-silico* data obtained regarding mitochondria machinery association with DHQ, we believed that the compound might cause ROS generation in *T. gondii* tachyzoites and eventually mitochondrial membrane disruption. All these predictions will need further investigation using *in vitro* experiments.

## Materials and Methods

### Molecular Docking of DHQ

The protein-ligand docking approach was employed to analyze and identify the specific amino acid interactions between DHQ and the respective receptors found on *T. gondii*. The binding affinities afford a good prediction of the ability of DHQ to inhibit *T. gondii* via various mechanisms modulated by proteins/enzymes (DNA gyrase, Calcium Dependent Protein Kinase 1 (CDPK 1), and prolyl tRNA synthetase) understudy.

The following platforms were used for the computational studies of the effect of DHQ on various replicative, and translational machinery; Vina Autodock, Pymol (2) (Schrodinger, LLC), PyRx, and Discovery Studio 2021.

### TgDNA gyrase structure modeling

Currently, the TgDNA gyrase crystal structure is not available. Therefore, an *in silico* model was made using the TgDNA sequence. In summary, the TgDNA gyrase subunit B sequence (ID KFG61327.1) was downloaded from the NCBI website. The sequence was converted into FASTA format using the sequence format converter (http://avermitilis.ls.kitasato-u.ac.jp/readseq.cgi), afterward, the sequence was modeled into its predicted 3D structure using the online SWISS-MODEL tool (https://swissmodel.expasy.org/). The model with the highest score in Global Model Quality Estimation (GQME) was utilized for the binding study.

### Ligand preparation

The DHQ was prepared as a ligand for docking onto the following receptors, 4DH4, ROP5C, ROP5B, and enzymes (TgDNA gyrase, Calcium Dependent Protein Kinase 1 (CDPK 1), and prolyl tRNA synthetase). These receptors/enzymes were selected for the molecular docking analysis to confirm the *in vitro* mechanism of action(s) based on their association with replication, invasion, and survival of *T. gondii*. Additionally, previous works on *Plasmodium spp* implicated dihydroquinine to inhibit DNA, RNA, and protein synthesis (Brossi et al., 1973; Brossi et al., 1971; Polet and Barr, 1968). The chemical structure of DHQ was extracted from the PubChem database. The ligand was uploaded into PyRx software via the Open babel plugin, and the 3D and geometry optimizations with energy minimization of DHQ structure were carried out. DHQ structure was converted to autodock DHQ.pdbqt ligand file via the same Open babel plugin in PyRx. The Vina wizard was used to load the DHQ.pdbqt file as a ligand to the respective receptors/enzymes used in the docking process. Huang et al. (2015), demonstrated that 5-aminopyrazole-4-carboxamide derivatives were selectively potent inhibitors of 4YJN (Calcium-Dependent Protein Kinase 1 (TgCDPK1) from *T. gondii*. They also showed the effectiveness of the structure-activity relationship of this compound in *a* mouse model against *T. gondii* in the brain, spleen, and peritoneal fluids. In this computational investigation, we analyzed the binding affinity of 5-aminopyrazole-4-carboxamide also to 4YJN to serve as an experimental control for all the binding studies of DHQ on the various molecules. The 2D structure (PDB format) of 5-aminopyrazole-4-carboxamide was downloaded from the PubChem database (CID_79254) and processed for docking. 5-aminopyrazole-4-carboxamide was prepared for docking following the same process as started earlier.

### Preparation of protein structures and grid generation

The Crystal structures of virulent alleles ROP5B (3Q5Z) (Reese and Boothroyd, 2011; Reese et al., 2014), and ROP5C (4VL8) (Reese et al., 2014) of *T. gondii* were downloaded from the RCSB homepage (http://www.rscb.org) at a resolution of 1.90 Å (Reese and Boothroyd, 2011), and 1.72 Å, respectively. Also, 4DH4 (Sommerville et al., 2013), 4YJN (Cardew et al., 2018), and 6A88 (Mishra et al., 2019) molecules were extracted from the same website at resolutions of 1.82 Å, 2.60 Å, and 2.60 Å, respectively. These proteins were loaded into Pymol 2 (Schrodinger, LLC), for detailed analysis of proteins’ structural elements, then each molecule was exported in a PDB file format with the removal of water molecules. Vina Autodock was used to adjust charges, check, and replace missing atoms and add polar hydrogens to the proteins structures. Next, all protein structures were energy minimized using the Autodock Vina. Additionally, proteins were loaded into PyRx and converted to receptors, the receptor grid boxes were generated in PyRx using the built-in Vina Wizard module, boxes were maximized to cover all active sites of the receptors. Docking of compounds to their respective receptors was achieved using the AutoDock tool in PyRx and analysis of specific amino acids in the binding pocket with DHQ or other ligands were achieved using Discovery Studio 2021.

## AUTHOR CONTRIBUTIONS

DAA conceived the idea. JAA performed the *in-silico* studies. JAA analyzed the data and drafted the first manuscript under the supervision of DAA. DAA, AN, and BK corrected the original manuscript. All authors approved the final manuscript.

## ACKNOWLEDGMENTS

None

## Supplementary Material

Supplementary Materials have been included in the main manuscript.

## Data Availability Statement

The datasets generated for this study are available upon request from the corresponding author.

